# Social selection maintains honesty of a dynamic visual signal in cichlid fish

**DOI:** 10.1101/039552

**Authors:** Judith C. Bachmann, Fabio Cortesi, Matthew D. Hall, N. Justin Marshall, Walter Salzburger, Hugo F. Gante

**Author notes:** Current address: Institute of Evolutionary Biology and Environmental Studies, University of Zurich, 8057 Zurich, Switzerland (Judith C. Bachmann).

## Abstract

How honest signals evolve is a question that has been hotly debated by animal communication theoreticians and for which empirical evidence has been difficult to obtain. Theory predicts that, due to strong conflicts of interest, communication in aggressive contexts should be under strong selection for clear and reliable signaling. On the other hand, context-dependent signaling increases cheating opportunities, depending on how senders and receivers use, acquire and process signal information. Using animal signaling theory, theoretical visual models and behavioral experimentation, we characterize and determine proximate honesty mechanisms of the facial coloration in the Princess of Burundi cichlid, Neolamprologus brichardi, a species with complex social interactions. We show that this facial color pattern evolved stable chromatic conspicuousness for efficient transmission in the aquatic environment, while context-dependent plasticity in luminance of the horizontal black stripe element is used to signal switches in aggressive intent. Importantly, using experimental signal manipulation we demonstrate that social selection by receiver retaliation is the mechanism responsible for maintaining signal honesty. We suggest that by affecting the evolution of pigmentation patterns in sexually monochromatic cichlid species, social selection can have potential impacts on diversification dynamics.

## INTRODUCTION

Our understanding of animal communication has been driven by advances in theory, not least because empirical evidence has been difficult to obtain [1]. Costly signaling theory is the dominant paradigm explaining the evolution of honest communication [2-5]. According to it, honesty is maintained by imposing different strategic costs on signals produced by animals of different qualities (e.g. handicaps and indices) [6-8]. In spite of generalized acceptance, other models have suggested that strategic costs at equilibrium are alone not sufficient, nor even necessary, for reliable signaling [9-11]. One alternative solution to the puzzling evolution of honest signals comes from potential, rather than realized costs, imposed by receivers. Indeed several theoretical models suggest that honesty of signaling systems can instead be socially-enforced and context-dependent. In this case animals that signal reliably do not need to incur any strategic costs on top of the efficacy costs that signal transmission entails [12-14]. These conclusions require that empiricists measure marginal costs of cheating in manipulated out-of-equilibrium signals where individuals are forced to exhibit unreliable signal expression [13, 15-18]. Here we quantify the signaling efficacy and message of the facial color pattern in the Princess of Burundi cichlid *(Neolamprologus brichardi)* using theoretical visual models and staged dyadic combats. By manipulating signal expression and simulating a cheater invasion we demonstrate that social selection promotes the honesty of this dynamic conventional signal with low production costs. By directly probing the sender of a signal, social selection is likely to be the mechanism of choice shaping the evolution of cheap context-dependent signals. In the same way that sexual selection drives the evolution of coloration in dichromatic species [19], we suggest that social selection can affect the evolution of pigmentation patterns in sexually monochromatic cichlid species, with potential impacts on diversification dynamics.

## RESULTS AND DISCUSSION

### A new framework for studying intraspecific color signals

The understanding of honest signaling in animal communication centers on the costs of expressing a signal, yet it remains unclear whether signaling costs have even been determined empirically (e.g. [17, 18, 20]). Here we combine conceptual approaches from visual modeling and signaling theory (e.g. [21]) into a new 3-stage framework that generates objective predictions about the evolution of reliable color signals, making the demonstration of the existence of strategic costs a more tractable empirical problem (Figure 1). We follow Higham’s [17] definition of costly signaling, where cost functions can be zero at the equilibrium, to include social selection through punishment as a mechanism that can generate marginal costs to cheaters and maintain signaling reliability (as elaborated elsewhere [20]). Our approach (Figure 1), which can be readily extended to other species, first requires using visual models to formally quantify signal efficacy and identify the correct target of communication. We then use behavior observations and assays to determine the message conveyed by our signal of interest. Finally, we identify which class of costs unreliable signaling might incur, by experimentally manipulating sender signals out-of-equilibrium and recording receiver’s reactions. Here we follow Fraser’s [20] classification based on intrinsic and imposed costs to determine whether liar detection mechanisms exist. Liar detection is expected to evolve in cheap conventional signals where receivers can immediately probe senders, but not in intrinsically costly handicaps or indices in which case reliability is verified far into the future in terms of viability and fecundity [13].

**Figure 1.**
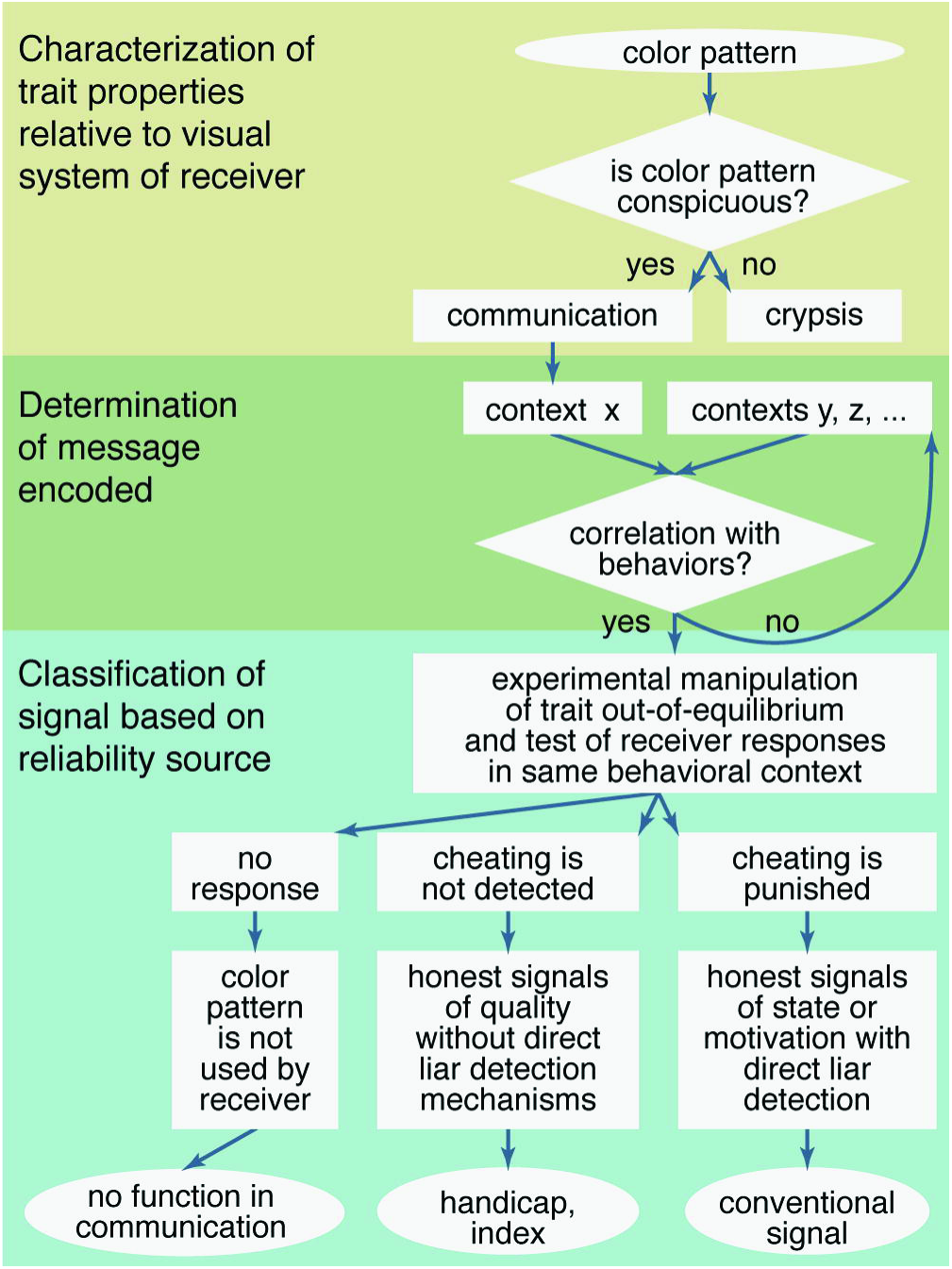
Proposed framework for studying color signals. Flowchart of the conceptual framework proposed for studying intraspecific color signals generates predictions to determine signal efficacy, function and proximate reliability mechanisms.

Using this framework we explored the evolution of the facial color mask in the cichlid fish *N. brichardi* (Figure 2). We chose this system because it is a lifelong territorial species with elaborate social habits for which considerable behavioral and ecological information is available (see Supplemental Experimental Procedures). It is a sexually monochromatic (i.e. both sexes look alike) substrate brooder of the species-rich tribe Lamprologini from East African Lake Tanganyika [22] and has emerged as a model in cooperative breeding studies [23]. The dominant couple has the peculiarity of being aided by several subordinate helpers in these tasks, organized in a linear hierarchy. Their rocky territory is a valuable resource that simultaneously provides substrate for reproduction and shelter against predation. Hence, losing access to a shelter has a strong negative impact on fitness and survival. As a consequence of cooperative breeding and colony life, individuals repeatedly and regularly interact, which creates optimal conditions for the evolution of context-dependent signaling by individuals of both sexes and different ages throughout their lives.

**Figure 2.**
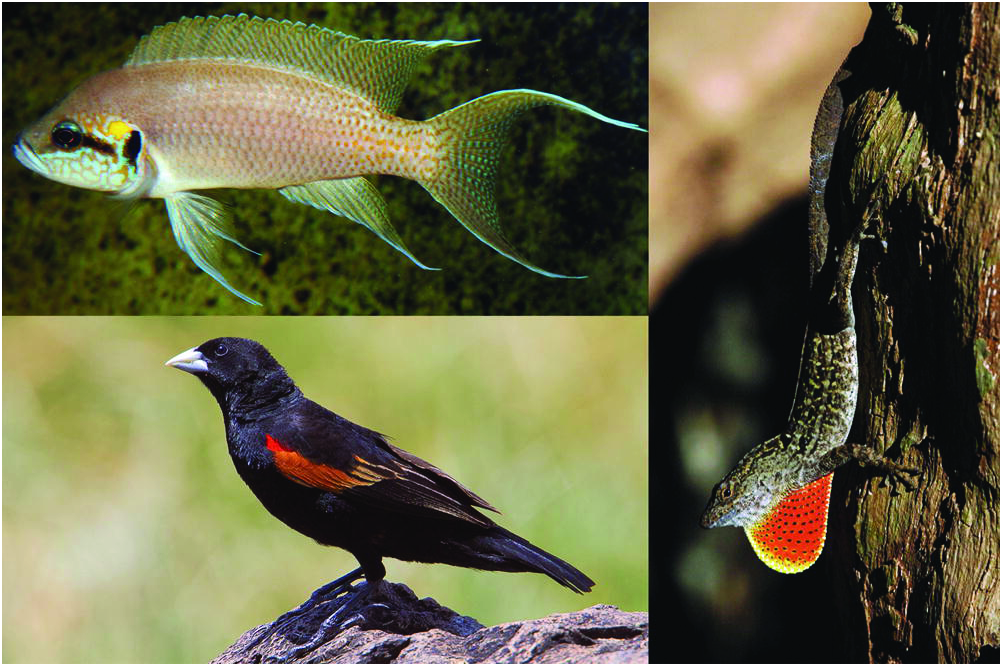
Dynamic animal visual signals. Territorial species display a variety of conspicuous visual signals to communicate aggressive intent. To decrease predation pressure and in non-aggressive contexts several species use morphological, physiological or behavioral adaptations to conceal signals [26, 44, 45, 52]. We propose that rapid physiological color changes, achieved by pigment movement in melanophores (black pigment cells), are a cheap proximate mechanism turning a visual signal of aggressive intent ‘on’ or ‘off’ in lifelong territorial fish. Clockwise from top left: facial color pattern in dominant Princess of Burundi cichlid *(Neolamprologus brichardi)*; extended dewlap in trunk-ground Brown Anole *(Anolis sagrei)*; partially covered epaulette in Fan-tailed Widowbird *(Euplectes axillaris)*.

### Stage 1: High chromatic conspicuousness of *N. brichardi*’s facial coloration

Unambiguous communication selects for signaling systems that promote effective stimulation of sensory systems relative to environmental noise and signal degradation. Such high conspicuousness to intended receivers is achieved by stimulation of adjacent photoreceptors in opposite ways by complementary radiance spectra [24-26]. Design strategies for increased conspicuousness and transmission efficacy thus include the use of (*i*) white or highly reflective colors adjacent to dark patches, (*ii*) adjacent patches with complementary colors and (*iii*) color combinations centered or just offset transmission maximum of the medium [24, 26, 27]. Further, a visual signal in a particular light environment is most conspicuous when adjacent color elements have greater contrasts than non-adjacent elements [27-30].

Using spectral reflectance measurements and theoretical fish visual models, we show that the facial color pattern in aggressive, dominant *N. brichardi* achieves high chromatic conspicuousness to the visual system of conspecifics by following all three predictions (Figure 3A and 3C, filled circles; Figure S1). This signal design is exceptionally effective and ensures transmission efficacy in the aquatic environment: white is a broadband optical reflector, reflecting across all the available light spectrum and structural blue patches reflect the high-intensity wavelengths available underwater, while the adjacent black melanic stripes absorb most incident light. Chromatic contrast is further achieved by use of complementary colors, blue and yellow, centered in the highest light intensity of water transmission. As such, chromatic contrasts differ significantly between adjacent and non-adjacent patches (linear mixed-effects model [LMM]: *F*_1,9_ = 207.31, P < 0.001) and all pairwise color comparisons are well above the just noticeable difference (JND) threshold of one, a standard in chromatic color discrimination [29, 31]; Figure 3A and 3C, filled circles). Compared to chromatic contrasts, achromatic contrasts do not seem to contribute to pattern conspicuousness, as adjacent and non-adjacent elements do not significantly differ in luminance from one another (LMM: *F*_1,9_ = 4.61, P = 0.06; Figure 3D, filled circles).

**Figure 3.**
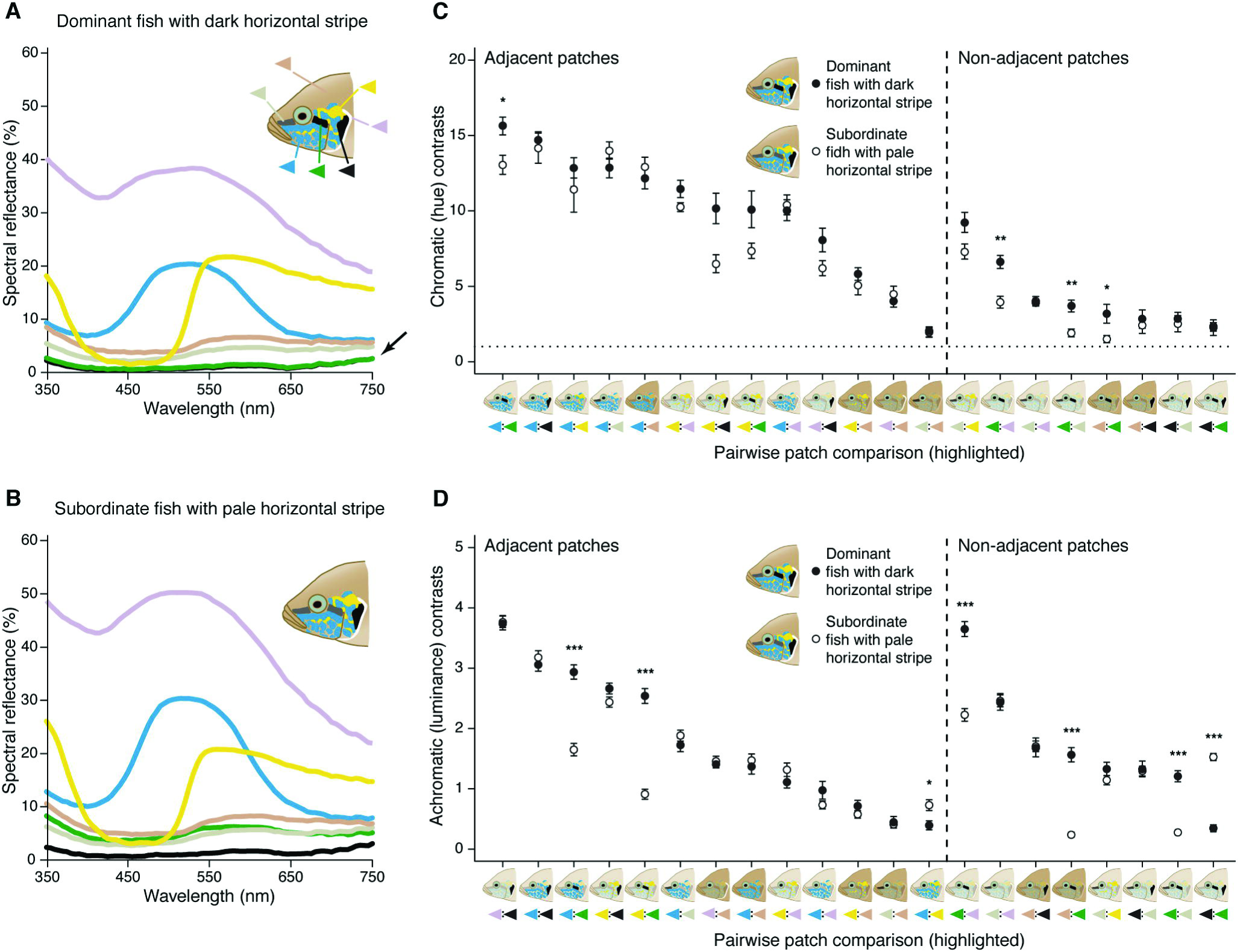
Color properties of facial elements in dominant and subordinate *Neolamprologus brichardi*. (A and B) Mean spectral reflectance of facial color patches. (A) Horizontal (green triangle) and vertical (black triangle) facial stripes have the same reflectance in dominant fish (note arrow). (B) Losing a combat and becoming subordinate significantly increases reflectance of horizontal facial stripe in subordinate fish, i.e. paling occurs. (See Figure S1G for 95% confidence intervals of spectral reflectance). (C and D) Chromatic and achromatic contrasts (mean ± SEM) between pairs of adjacent and non-adjacent color patches as perceived by *N. brichardi*, ordered from highest to lowest in dominant fish. (C) High chromatic contrast Δ*S* is achieved by any combination of blue, yellow and black patches. (D) High achromatic contrast Δ*L* is achieved by combining black melanic stripes and other patches. Stippled line marks the 1 JND (just noticeable difference), threshold after which two colors are thought to be perceived as different [29, 53]. Asterisks illustrate significant differences in contrast between dominant and subordinate fish (*** P < 0.001, ** P < 0.01, * P < 0.05). (Figure S1 shows data used to build visual models)

### Stage 2: *Neolamprologus brichardi* make context-dependent use of facial signal

High chromatic conspicuousness of facial patterns implicates selection for unambiguous signaling, at least at close range (Figure 1). We thus tested its function in communication by staging dyadic combats of territory-holding fish. As expected, body size (LMM: *F*_1,18_ = 8.02, *P* = 0.01) and fighting ability (LMM: *F*_1,18_ = 67.31, *P* < 0.001) determine the outcome of staged combats, irrespective of sex (LMM: *F*_1,18_ = 1.85, *P* = 0.19 and LMM: *F*_1,18_ = 0.04, *P* = 0.85; Figure S2A and S2B). Most importantly, we found that a change in aggressive intent by losers of the combat leads to a rapid paling of the horizontal facial stripe at the end of the contest (generalized linear mixed-effects model [GLMM] with binomial error distribution: 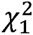 = 14.97, *P* < 0.001; Figures 3B, 4A, S1G). Hence stripe intensity at the end of the combat reflects motivation to fight and aggressive intent, while stripe darkness at the beginning does not influence contest outcome (GLMM with binomial error distribution: 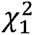 = 0.01, P = 0.93), which is fundamentally different from other well-described signals that function as badges of status [31]. Therefore, rapid paling of the horizontal facial stripe may be used to instantaneously signal an individual’s intent to fight and dominance. Such rapid movement of pigments within melanophores (black pigment cells) is a physiological response available to many lower vertebrates (e.g. fish, reptiles) and invertebrates (e.g. cephalopods), and can occur within a few seconds in fish [32, 33].

**Figure 4.**
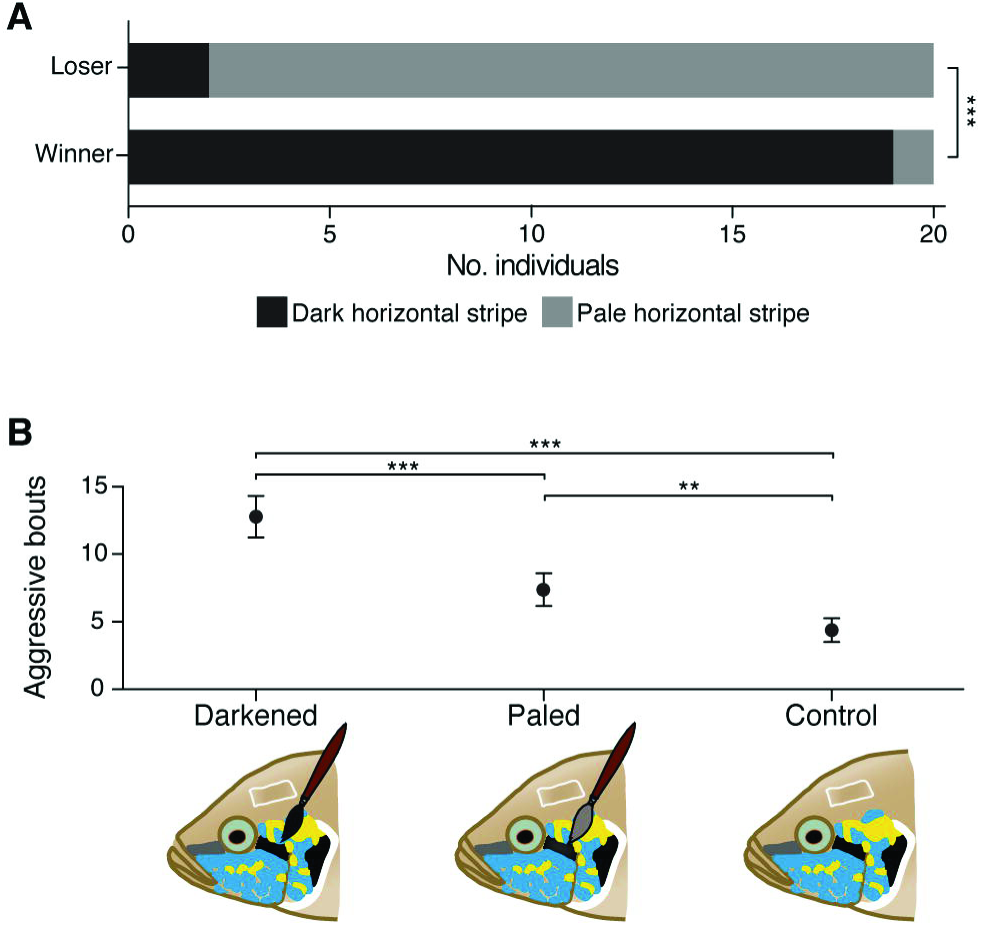
Horizontal facial stripe provides reliable information on aggressive intent. (A) Facial stripe intensity is associated with fighting ability (winning or losing) at end of combat. L: losers; W: winners. (B) Social selection (aggressive bouts, mean ± SEM) on out-of-equilibrium signals and control. Unreliable signaling of strength (darkened stripe) and weakness (paled stripe) are punished by increased receiver retaliation costs relative to reliable signaling (control). Asterisks illustrate significant differences in facial stripe luminance at end of combat and of pairwise post-hoc tests between treatments (*** P < 0.001, ** P < 0.01). (See also Figure S2).

Next, we used theoretical visual models to test whether the physiological paling of the horizontal stripe induces changes in conspicuousness of the overall facial pattern. We found that even after paling takes place chromatic conspicuousness is unaffected (empty circles in Figures 3C and S1H). In particular, high chromatic contrast is achieved by adjacent and non-adjacent signal design (LMM: *F*_1,18_ = 208.21, *P* < 0.001) and not by stripe darkness (LMM: *F*_1,18_ = 3.48, *P* = 0.08) or interaction between the two fixed effects (LMM: *F*_1,18_ = 0.05, *P* = 0.82). This model explains 99.31% of chromatic contrast variance, 96.50% of which is explained by adjacency of the color elements, while changes in horizontal stripe luminance explain the remaining variance. On the other hand, we found that achromatic contrasts are strongly influenced by changes in luminance of the horizontal stripe (LMM: *F*_1,18_ = 9.11, *P* = 0.007), as the balance between adjacent and non-adjacent contrasts (LMM: *F*_1,18_ = 5.07, *P* = 0.037) and the interaction between the two becomes important (LMM: *F*_1,18_ = 6.78, *P* = 0.018; empty circles in Figure 3D and S1I). This model explains 95.90% of the achromatic contrast variance, 68.53% of which is explained by changes in darkness of the horizontal stripe, 22.34% by signal design (patch adjacency) and the remainder 5.02% by their interaction. Thus, we find that white, yellow and blue are less dynamic elements of the facial color pattern, and seem to provide little or no information regarding changes in aggressive intent but instead act as amplifiers to enhance pattern conspicuousness.

Our visual models and behavioral experiments indicate that individuals use rapid physiological changes in luminance (achromatic contrast) of the horizontal stripe element to dynamically communicate reversals in aggressive intent and dominance, while the color pattern conspicuousness remains high at all times by virtue of its stable chromatic properties. Using this dual mechanism is an elegant way to ensure that conspicuousness, and hence communication efficacy, does not decrease due to context-dependent signaling. This constantly ‘on’ signaling strategy of aggressive intent in *N. brichardi* is unexpected as it is opposite to other signaling systems such as in anoles lizards, chameleons, or transiently territorial fish which only briefly or seasonally display their signals during agonistic encounters [32, 34-36] (Figure 2). Our findings could possibly be explained by lifelong territoriality, constant interactions with conspecifics and different predation escape strategies of *N. brichardi*. While these stenotopic cichlids rely on their valuable rocky territories for shelter (and breeding) and conspicuously signal their ownership at all times, chameleons have to rely on immobility and camouflage to escape avian predation and become only momentarily conspicuous while displaying to conspecifics [35]. Instead, from a signaling perspective uninterrupted conspicuousness of *N. brichardi* is more similar to that of aposematic species, which rely on high conspicuousness to continuously signal their distastefulness [37, 38].

### Stage 3: Proximate mechanisms producing an evolutionary stable signaling strategy

Using a dyadic combat experiment in combination with the visual models we showed that changes in luminance of the horizontal melanistic stripe are used during social agonistic interactions and correlate with aggressive intent. However, whether the fish directly respond to the physiological color changes of the horizontal facial stripe and if these changes are then used to assess another individual’s aggressive intentions needs direct behavioral evaluation. Moreover, if changes in luminance were to reliably signal aggressive intent, we would expect the existence of an honesty mechanism to minimize cheating opportunities. To test these expectations we simulated a cheater invasion of the signaling system, by manipulating luminance of the horizontal facial stripe out-of-equilibrium (via artificial darkening or paling; Figures S2C, S2D and Table S1) and presenting fish to their mirror images. Our setup is opposite to the commonly used approach of displaying manipulated individuals to focal territory owners as noted by Bradbury and Vehrencamp [3], having the advantage of testing behavior of non-territorials (i.e. the receivers of the mirror image), which are the ones most interested in detecting unreliable signals if used by territorial, dominant individuals. As a null hypothesis (Figure 1), (1) we do not expect to observe differences in aggression toward manipulated or non-manipulated individuals if stripe intensity does not encode individual fighting abilities (but simply correlates with them). On the other hand, (2) if stripe intensity signals a contest-independent intrinsic quality whereby strategic costs guarantee honesty (e.g. handicap), subordinates should not challenge cheating individuals with enhanced signals but should do so toward individuals with subdued signals. Alternatively, (3) if stripe intensity signals contest-dependent dominance whereby social costs (i.e. punishment of cheaters) maintain signal honesty, we expect increased levels of aggression toward any unreliable signal (i.e. a conventional signal).

We found that receivers actively ‘read’ and react to manipulations of the horizontal facial stripe, recognizing and punishing cheaters (Figures 4B and S2E-G). Manipulation of the horizontal stripe had a significant effect on the number of aggressive bouts received (LMM: *F*_2,45_ = 13.73, *P* < 0.001), irrespective of sex (LMM: *F*_2,45_ = 0.48, *P* = 0.62). Individuals with darkened stripes received significantly more aggression than individuals with paled stripes (Tukey HSD: *z* = −3.89, *P* < 0.001) and controls (Tukey HSD: *z* = −6.59, *P* < 0.001). Importantly, individuals with paled stripes also received more aggression than controls (Tukey HSD: *z* = −2.97, *P* = 0.008), indicating that unreliable signaling brings increased marginal costs to all types of cheaters.

By manipulating the signal out-of-equilibrium we simultaneously show that physiological color changes are interpreted by receivers as a dynamic context-dependent signal of aggressive intent and that social selection by receiver retaliation is the proximate mechanism effectively promoting the honesty of this visual signal (hypothesis 3, above). Thus, as with communication efficacy, we demonstrate that communication reliability does not decrease due to context-dependent signaling but is rather promoted by contest-dependent policing. Since aggressive intent is not a quality that can be easily handicapped [3], receivers can directly assess reliability of signals of aggressive intent with relative ease [13] and impose social costs on cheaters. Our study provides rare empirical evidence that, similar to paper wasps [31, 39], fish are able to detect and punish individuals who signal unreliably, be they cheaters signaling strength (‘bluffers’) or modest liars (‘Trojans’). Interestingly, the fact that social selection against cheaters is not symmetrical supports the view that signaling systems are more likely disrupted by ‘Trojans’ than by ‘bluffers’ [40]. We thus provide empirical support to theoretical models concluding that honest communication does not require signals with differential strategic costs and that reliability can indeed be guaranteed by mechanisms that promote low realized costs for honest signalers, such as social selection [12-14, 16]. Hence, since receivers can effectively probe reliability of signals in real time, we propose that social selection and cheap conventional signals are expected to be a widely chosen solution for honest context-dependent signaling.

Physiological color changes have previously been implicated in signaling aggressive intent in a number of taxa, in particular fish [32, 41-43]. Increased levels of aggression toward the signal reported in some of these studies were interpreted as receiver retaliation costs maintaining honesty of a conventional signal. We extend these findings and show that unreliable signaling has increased costs relative to reliable signaling, which is pivotal to the evolution of honest signals [2, 13, 17]. The rapid physiological color changes of this dynamic color signal rival the morphological and behavioral context-dependent signaling strategies evolved by other territorial species [44, 45] and allow the fish to instantaneously communicate their intention to fight or retreat from the combat by reliably showing or concealing the signal of aggressive intent. Such dynamic expression noted among fish [46] contrasts with other more or less static visual signals, such as plumage badges of status in birds [47, 48], exoskeleton color patterns in insects [31], or aposematic signals in poison frogs [37], all of which are thought to signal more temporally stable characteristics of quality and distastefulness.

In summary, our framework for color signal analysis proved important in generating predictions according to trends emerging from theory. We demonstrated that the facial mask of *N. brichardi* has stable chromatic properties that keep signaling efficacy high at all times, while rapid physiological changes in luminance of just one element (the horizontal melanic stripe) communicate reversals in aggressive intent and dominance. We further demonstrated that social selection maintains honesty of the signaling system, which could be nature’s favored mechanism for promoting the honesty of dynamic visual signals with low production costs, such as those produced by physiological color changes. Together, these findings suggest that social selection may account for the dramatic diversity of color patterns (stripes, bars, blotches) we observe in many sexually monochromatic cichlid species endemic to Lake Tanganyika [22] and elsewhere, acting together with natural selection in shaping diversity in cichlid fishes [49, 50]. Social selection is expected to drive rapid signal evolution especially in isolated allopatric populations [14, 51], but until now research into color signaling in cichlids has centered on the sexually dichromatic assemblages from Lake Malawi and Lake Victoria [19]. Our results point to rapid social trait evolution as another process potentially affecting speciation dynamics in cichlids. Confirmation of its importance would bring social selection to the same level as sexual and natural selection in shaping adaptive radiations of cichlid fishes.

## AUTHOR CONTRIBUTIONS

H.F.G. conceived the study and designed the experiments together with J.C.B., F.C. and W.S., J.C.B., F.C. and H.F.G. performed the experiments and analyzed the data together with M.D.H. All authors contributed to writing and discussion. All authors reviewed and approved the final version of the manuscript.

## ACKNOWLEDGMENTS

This work was supported by the Swiss National Science Foundation (SNSF) grant 138224 to WS. Photographs of the anole and widowbird are a courtesy of Melisa Losos and Jan Willem Steffelaar. We thank designers Inês Santiago and Marco Silva for comments on figure preparation, and Adrian Indermaur for fishkeeping.

## REFERENCES

1. Grose, J. (2011). Modelling and the fall and rise of the handicap principle. Biol. Philos. 26, 677–696.

2. Searcy, W. A., and Nowicki, S. (2005). The evolution of animal communication: reliability and deception in signaling systems (Princeton: Princeton University Press).

3. Bradbury, J. W., and Vehrencamp, S. L. (2011). Principles of Animal Communication, Second Edition (Sinauer Associates, Inc.; 2 edition).

4. John Maynard-Smith, and Harper, D. (2003). Animal Signals First edit. (Oxford: Oxford University Press).

5. Hurd, P. L., and Enquist, M. (2005). A strategic taxonomy of biological communication. Anim. Behav. 70, 1155–1170.

6. Zahavi, A. (1975). Mate selection-A selection for a handicap. J. Theor. Biol. 53, 205–214.

7. Biernaskie, J. M., Grafen, A., and Perry, J. C. (2014). The evolution of index signals to avoid the cost of dishonesty. Proc. R. Soc. B Biol. Sci. 281.

8. Grafen, A. (1990). Biological signals as handicaps. J. Theor. Biol. 144, 517–546.

9. Getty, T. (1998). Reliable signalling need not be a handicap. Anim. Behav. 56, 253–255.

10. Getty, T. (1998). Handicap signalling: when fecundity and viability do not add up. Anim. Behav. 56, 127–130.

11. Hurd, P. L. (1997). Is signalling of fighting ability costlier for weaker individuals? J. Theor. Biol. 184, 83–88.

12. Hurd, P. L. (1995). Communication in discrete action-response games. J. Theor. Biol. 174, 217–222.

13. Lachmann, M., Számadó, S., and Bergstrom, C. T. (2001). Cost and conflict in animal signals and human language. Proc. Natl. Acad. Sci. U. S. A. 98, 13189–13194.

14. Tanaka, Y. (1996). Social selection and the evolution of animal signals. Evolution (N. Y). 50, 512–523.

15. Kotiaho, J. S. (2001). Costs of sexual traits: a mismatch between theoretical considerations and empirical evidence. Biol. Rev. Camb. Philos. Soc. 76, 365–76.

16. Számadó, S. (2011). The cost of honesty and the fallacy of the handicap principle. Anim. Behav. 81, 3–10.

17. Higham, J. P. (2014). How does honest costly signaling work? Behav. Ecol. 25, 8–11.

18. Számadó, S. (2012). The rise and fall of handicap principle: a commentary on the “Modelling and the fall and rise of the handicap principle.” Biol. Philos. 27, 279–286.

19. Wagner, C. E., Harmon, L. J., and Seehausen, O. (2012). Ecological opportunity and sexual selection together predict adaptive radiation. Nature 487, 366–9.

20. Fraser, B. (2011). Costly signalling theories: beyond the handicap principle. Biol. Philos. 27, 263–278.

21. Laidre, M. E., and Johnstone, R. A. (2013). Animal signals. Curr. Biol. 23, R829–R833.

22. Gante, H. F., and Salzburger, W. (2012). Evolution: cichlid models on the runaway to speciation. Curr. Biol. 22, R956–958.

23. Wong, M., and Balshine, S. (2011). The evolution of cooperative breeding in the African cichlid fish, Neolamprologus pulcher. Biol. Rev. Camb. Philos. Soc. 86, 511–530.

24. Lythgoe, J. N. (1979). The ecology of vision (Oxford: Clarendon Press).

25. Hurvich, L. M. (1981). Colour vision (Sinauer Associates Inc., U.S.).

26. Marshall, N. J. (2000). Communication and camouflage with the same “bright” colours in reef fishes. Philos. Trans. R. Soc. Lond. B. Biol. Sci. 355, 1243–1248.

27. Endler, J. A. (1992). Signals, signal conditions, and the direction of evolution. Am. Nat. 139, S125–S153.

28. Endler, J. A. (1990). On the measurement and classification of colour in studies of animal colour patterns. Biol. J. Linn. Soc. 41, 315–352.

29. Endler, J. A. (2012). A framework for analysing colour pattern geometry: adjacent colours. Biol. J. Linn. Soc. 107, 233–253.

30. Guilford, T., and Dawkins, M. S. (1993). Receiver psychology and the design of animal signals. Trends Neurosci. 16, 430–436.

31. Tibbetts, E. A., and Dale, J. (2004). A socially enforced signal of quality in a paper wasp. Nature 432, 218–222.

32. Muske, L. E., and Fernald, R. D. (1987). Control of a teleost social signal. I. Neural basis for differential expression of a color pattern. J. Comp. Physiol. A 160, 89–97.

33. Fujii, R. (2000). The regulation of motile activity in fish chromatophores. Pigment Cell Res. 13, 300–319.

34. Leal, M., and Fleishman, L. J. (2004). Differences in visual signal design and detectability between allopatric populations of anolis lizards. Am. Nat. 163, 26–39.

35. Stuart-Fox, D., and Moussalli, A. (2008). Selection for social signalling drives the evolution of chameleon colour change. PLoS Biol. 6, e25.

36. Losos, J. B. (2009). Lizards in an evolutionary tree: ecology and adaptive radiation of anoles (Berkeley: University of California Press).

37. Darst, C. R., Cummings, M. E., and Cannatella, D. C. (2006). A mechanism for diversity in warning signals: conspicuousness versus toxicity in poison frogs. Proc. Natl. Acad. Sci. U. S. A. 103, 5852–5857.

38. Cheney, K. L., Cortesi, F., How, M. J., Wilson, N. G., Blomberg, S. P., Winters, A. E., Umanzör, S., and Marshall, N. J. (2014). Conspicuous visual signals do not coevolve with increased body size in marine sea slugs. J. Evol. Biol. 27, 676–687.

39. Tibbetts, E. A., and Izzo, A. (2010). Social punishment of dishonest signalers caused by mismatch between signal and behavior. Curr. Biol. 20, 1637–1640.

40. Számadó, S. (2011). Long-term commitment promotes honest status signalling. Anim. Behav. 82, 295–302.

41. Korzan, W., Summers, T., and Summers, C. (2002). Manipulation of visual sympathetic sign stimulus modifies social status and plasma catecholamines. Gen. Comp. Endocrinol. 128, 153–161.

42. Moretz, J. A., and Morris, M. R. (2003). Evolutionarily labile responses to a signal of aggressive intent. Proc. R. Soc. B Biol. Sci. 270, 2271–2277.

43. Rodrigues, R. R., Carvalho, L. N., Zuanon, J., and Del-Claro, K. (2009). Color changing and behavioral context in the Amazonian dwarf cichlid Apistogramma hippolytae (Perciformes). Neotrop. Ichthyol. 7, 641–646.

44. Hansen, A. J., and Rohwer, S. (1986). Coverable badges and resource defence in birds. Anim. Behav. 34, 69–76.

45. Metz, K. J., and Weatherhead, P. J. (1992). Seeing red: uncovering coverable badges in red-winged blackbirds. Anim. Behav. 43, 223–229.

46. Mäthger, L. M., Land, M. F., Siebeck, U. E., and Marshall, N. J. (2003). Rapid colour changes in multilayer reflecting stripes in the paradise whiptail, Pentapodus paradiseus. J. Exp. Biol. 206, 3607–3613.

47. Rohwer, S. (1975). The social significance of avian winter plumage variability. Evolution (N. Y). 29, 593–610.

48. Senar, J. C. (1999). Plumage coloration as a signal of social status. In Proceedings of the International Ornithological Congress, N. J. Adams and R. H. Slotow, eds. (Johannesburg: BirdLife South Africa), pp. 1669–1686.

49. Salzburger, W. (2009). The interaction of sexually and naturally selected traits in the adaptive radiations of cichlid fishes. Mol. Ecol. 18, 169–185.

50. Muschick, M., Indermaur, A., and Salzburger, W. (2012). Convergent evolution within an adaptive radiation of cichlid fishes. Curr. Biol. 22, 2362–2368.

51. West-Eberhard, M. J. (1983). Sexual selection, social competition, and speciation. Q. Rev. Biol. 58, 155–183.

52. Siebeck, U. E., Parker, A. N., Sprenger, D., Mäthger, L. M., and Wallis, G. (2010). A species of reef fish that uses ultraviolet patterns for covert face recognition. Curr. Biol. 20, 407–410.

53. Vorobyev, M., and Osorio, D. (1998). Receptor noise as a determinant of colour thresholds. Proc. Biol. Sci. 265, 351–358.

